# Widespread pleiotropy confounds causal relationships between complex traits and diseases inferred from Mendelian randomization

**DOI:** 10.1101/157552

**Authors:** Marie Verbanck, Chia-Yen Chen, Benjamin Neale, Ron Do

## Abstract

A fundamental assumption in inferring causality of an exposure on complex disease using Mendelian randomization (MR) is that the genetic variant used as the instrumental variable cannot have pleiotropic effects. Violation of this ‘no pleiotropy’ assumption can cause severe bias. Emerging evidence have supported a role for pleiotropy amongst disease-associated loci identified from GWA studies. However, the impact and extent of pleiotropy on MR is poorly understood. Here, we introduce a method called the Mendelian Randomization Pleiotropy RESidual Sum and Outlier (MR-PRESSO) test to detect and correct for pleiotropy in multi-instrument summary-level MR testing. We show using simulations that existing approaches are less sensitive to the detection of pleiotropy when it occurs in a subset of instrumental variables, as compared to MR-PRESSO. Next, we show that pleiotropy is widespread in MR, occurring in 41% amongst significant causal relationships (out of 4,250 MR tests total) from pairwise comparisons of 82 complex traits and diseases from summary level genome-wide association data. We demonstrate that pleiotropy causes distortion between-168% and 189% of the causal estimate in MR. Furthermore, pleiotropy induces false positive causal relationships-defined as those causal estimates that were no longer statistically significant in the pleiotropy corrected MR test but were previously significant in the naive MR test-in up to 10% of the MR tests using a *P* < 0.05 cutoff that is commonly used in MR studies. Finally, we show that MR-PRESSO can correct for distortion in the causal estimate in most cases. Our results demonstrate that pleiotropy is widespread and pervasive, and must be properly corrected for in order to maintain the validity of MR.

---

Epidemiological studies have established correlations among numerous exposures and complex diseases^1^. It can be challenging to draw causal inference from these studies due to reverse causation, confounding and/or other biases^2^.

Mendelian randomization (MR) is a commonly-used human genetics approach that can infer causality of an exposure with complex disease^3,4^. MR presents a number of advantages over observational epidemiology, including controlling for non-heritable environmental confounders in such analyses and the ability to use genetic instruments to evaluate the impact of an exposure without necessitating the collection of that exposure in the outcome group. MR utilizes genetic variants as genetic instrumental variables (IVs) that are robustly associated with the exposure of interest and tests whether these variants results in a proportional effect on the outcome.

With the discovery of thousands of trait-associated loci identified from genome-wide association (GWA) studies, multi-instrument MR methods have been developed to aggregate estimates from multiple IVs, building on the rich history of mediation analyses. The effect sizes of the variants on the exposure and the effect sizes on the outcome can be extracted from GWA summary statistics and multi-instrument MR can be performed in a linear regression framework^5,6^.

A fundamental assumption of MR is exclusion restriction which assumes that the IV used for MR analysis acts solely on the target outcome exclusively through the exposure of interest and does not have pleiotropy^2^. Pleiotropy occurs when the variant has an effect on other untested traits that have an effect on the target outcome, or a direct effect with the target disease outcome^7^. Specifically, violation of the exclusion restriction, or ‘no pleiotropy’ assumption can confound MR tests, leading to biased estimates and potentially, false positive causal relationships.

Emerging evidence have supported a pervasive role of pleiotropy amongst loci identified from GWA studies. Studies have shown that many traits are genetically correlated with each other^8^. Furthermore, studies have shown that hundreds of individual variants identified from GWA studies are associated with multiple traits^9-14^. Many of these variants for the most part have been used as IVs for causal inference in MR studies with the assumption that these variants do not have pleiotropic effects.

As a result, there has been recent discussion about whether pleiotropy may have serious consequences on the validity of MR analyses to date. Some have raised skepticism about the MR approach due to the pervasiveness of pleiotropy amongst trait-associated variants^15^, while others have defended MR by noting that pleiotropy has long been known to impose limits on MR^16^.

Despite these discussions, it is currently unknown the extent to which pleiotropy affects causal relationships inferred by MR. Importantly, an evaluation of pleiotropy and how it impacts the validity of MR testing across complex traits and diseases has not been performed in a systematic manner and at a large-scale.

Here, we developed the Mendelian Randomization Pleiotropy RESidual Sum and Outlier (MR-PRESSO) approach to detect and correct for pleiotropy in multi-instrument summary-level MR testing. We demonstrate the validity and sensitivity of MR-PRESSO using extensive simulations. Next, we leverage MR-PRESSO to evaluate the role of pleiotropy and strategies to correct for it, amongst >4,025 MR tests of complex traits and diseases derived from 82 summary level genome-wide association datasets.

## Results

MR-PRESSO is a unified framework that allows for the evaluation of pleiotropy in a standard MR model (see **Online Methods**, and **Supplementary Figures 1** and **2**). The method extends on previous approaches that utilize the general model of multi-instrument MR^17^ (see **Online Methods** and **Supplementary Methods**) and demonstrates better sensitivity and correction for pleiotropy as compared to previous approaches that use summary statistics from GWA in MR, including the MR-Egger intercept^18^ and the Q test^19^.

MR-PRESSO has three components (see Figure 1), including: 1) detection of pleiotropy (MR-PRESSO global test); 2) correction of pleiotropy via outlier removal (MR-PRESSO outlier test); and 3) testing of significant differences in the causal estimates before and after correction for outliers (MR-PRESSO distortion test). The MR-PRESSO global test evaluates overall pleiotropy amongst all IVs in a single Mr test by comparing the observed residual sum of squares (RSS) of the effect sizes of the variants on the outcome to the expected distribution of RSS. The MR-PRESSO outlier test extends on the global test by testing for the presence of variant effect sizes that are outliers by utilizing the observed and expected distributions of the specific tested variant. Finally, the MR-PRESSO distortion test evaluates the significance of the distortion between the causal estimate before and after removal of outliers (detected from the outlier test of MR-PRESSO) (see **Online Methods**).

**Figure 1.**
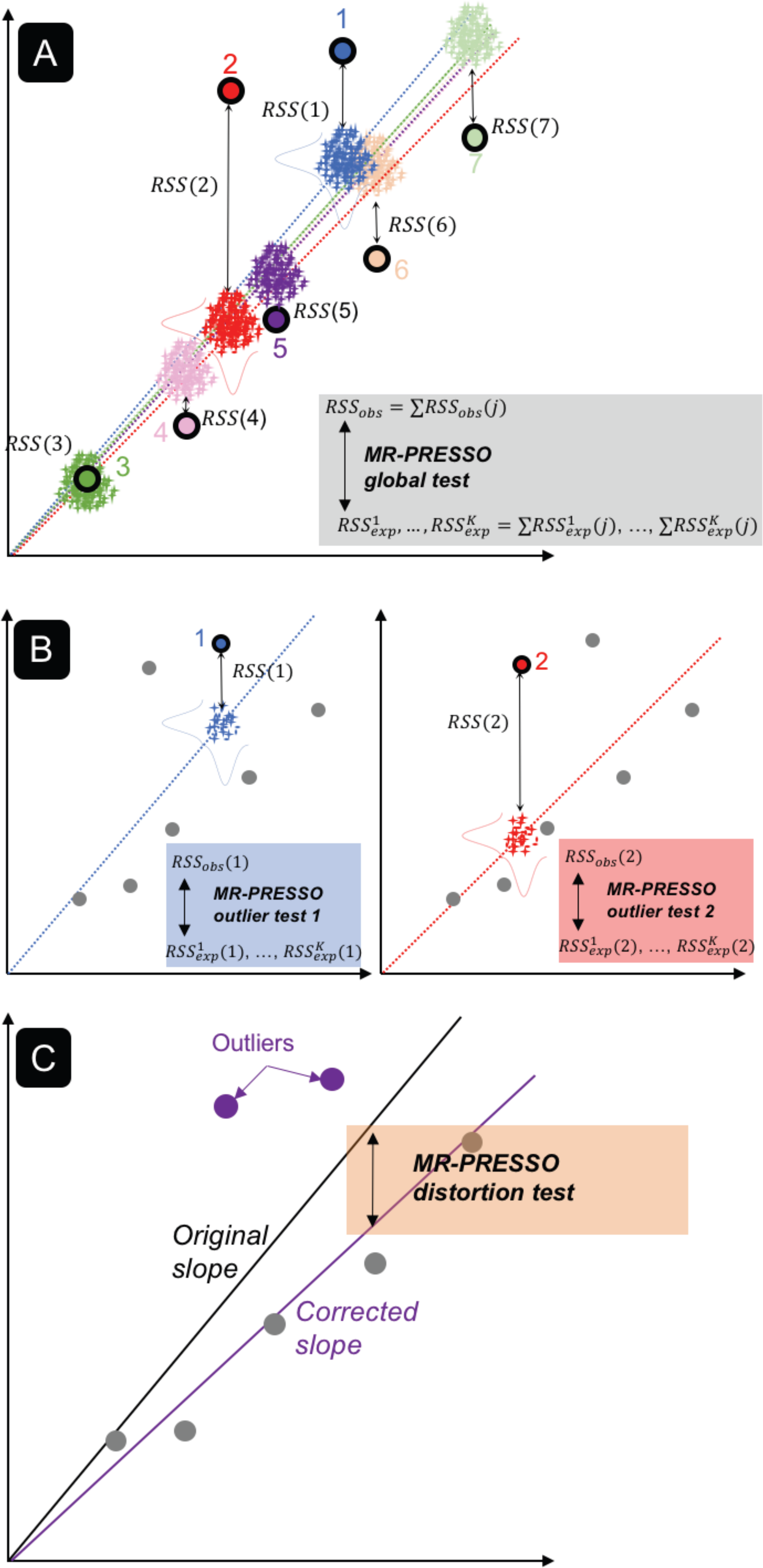
Description of the Mendelian Randomization Pleiotropy RESidual Sum and Outlier (MR-PRESSO) framework. The MR-PRESSO framework is comprised of three components. Panel A represents the global test. For each variant j, a slope is computed without the variant using standard IVW meta-analysis MR (colored dotted lines). The observed residual sum of squares RSS_obs_(j) is computed as the squared difference between the observed effect size of variant j on the outcome and the effect size predicted using the slope computed without j. In addition, K pairs of random effect sizes for the exposure (x-axis) and the outcome (y-axis) (represented as crosses) are drawn from two Gaussian distributions (horizontal and vertical bell curves respectively for the exposure and outcome) from the predicted effect sizes and standard errors using the slope computed without j. A distribution of K expected 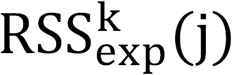 is then calculated. By summing up the J RSSobs(j), we have a global statistic that we can compare to the K expected sum of 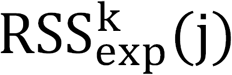 to compute an empirical *P-value.* Panel B represents the outlier test. The principle of the outlier test extends on the the principle of the global test and should be used if the global test is statistically significant. A test is performed for each variant j by comparing the observed RSS_obs_(j) to the K expected 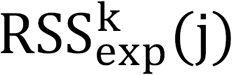, which allows to obtain an empirical *P-value* which is then multiplied by the number of variants J to account for multiple testing (Bonferroni correction). Here, only variants (1 and 2) are shown for simplicity. Panel C represents the distortion test. The panel shows how removing significant outliers detected by the MR-PRESSO outlier test (variant 1 and 2) leads to a noninflated slope used to estimate the causal effect. The distortion between the causal estimates can be evaluated using the MR-PRESSO distortion test.

### Comparison of statistical properties of methods to detect pleiotropy

We performed simulations to evaluate the statistical properties of MR-PRESSO and compared it to other existing methods, including the intercept test from MR-Egger, and the Q test (see **Online Methods**; **Supplementary Material**). For all three approaches, we performed 10,000 simulations under different scenarios to evaluate the statistical power (1-type II error) and false positive rate (type I error) for these methods. We simulated scenarios with varying parameters, including: 1) with (β_causal_ = 0.5) or without causal effects (β_causal_ = 0) from the exposure to the outcome; 2) with (β_pleiotropic_ = 0.5) or without (β_pleiotropic_ = 0) directional (all effect sizes in one direction) or balanced pleiotropy (approximately 50% of effect sizes in positive direction and 50% in the negative direction); and; 3) different proportion of variants with pleiotropy in the multi-instrument MR test (see **Online Methods**).

Our simulations showed that MR-PRESSO and the Q test were well powered when >1% of variants were under directional or balanced pleiotropy (power > 72%).

Importantly, the Q test produced an inflated false positive rate for pleiotropy detection (type I error = 39%), in the simulation with no pleiotropy and β_causal_ = 0.05 whereas MR-PRESSO showed no inflation (type I error = 5%) in the same simulation (Table 1). On the other hand, the intercept test from MR-Egger showed limited power to detect pleiotropy under all scenarios (power < 13%).

**Table 1.** Evaluation of statistical power (%) at a significance threshold of 0.05 for different methods to detect pleiotropy in Mendelian randomization.

| Causal<br>Effect $\beta_{causal}$ | Pleiotropy | Percentage<br>of Pleiotropy | Intercept test | Q test | MR-PRESSO |
| --- | --- | --- | --- | --- | --- |
|  |  |  | MR-Egger<br>regression | IVW meta-analysis | IVW meta-analysis |
| 0 | None | 0 | 4.96 | 5.3 | 5.4 |
| 0 | Directional | 1 | 4.87 | 24.4 | 24.4 |
| 0 | Directional | 5 | 6.65 | 85.9 | 86 |
| 0 | Directional | 10 | 10.85 | 99.4 | 99.3 |
| 0 | Balanced | 1 | 5.15 | 24.4 | 24.5 |
| 0 | Balanced | 5 | 4.78 | 87.5 | 87.4 |
| 0 | Balanced | 10 | 5.12 | 99.6 | 99.6 |
| 0.05 | None | 0 | 5.81 | 39 | 5.5 |
| 0.05 | Directional | 1 | 6.23 | 51.5 | 18 |
| 0.05 | Directional | 5 | 7.62 | 92.6 | 72.8 |
| 0.05 | Directional | 10 | 12.43 | 99.7 | 97.2 |
| 0.05 | Balanced | 1 | 5.49 | 51.5 | 17.4 |
| 0.05 | Balanced | 5 | 5.01 | 93.1 | 75.2 |
| 0.05 | Balanced | 10 | 5.26 | 99.7 | 98 |

variant minor allele frequency = 0.1; Number of variants = 100; sample size of GWAS for E1, E2 and Y = 1,000. InSIDE condition satisfied in all simulations.

Next, we compared the causal effect estimates using two different methods: Inverse Variance-Weighted (IVW) meta-analysis^17^ and MR-Egger regression^18^ (see **Online Methods; Supplementary Table 1**). As expected, the IVW meta-analysis displayed the most statistical power to detect a causal estimate among the three methods whereas MR-Egger regression had the lowest, as indicated by the smallest mean standard error (SE). The naive IVW meta-analysis produced biased effect estimates that increases with the proportion of variants with pleiotropy.

We next examined the impact of correcting for pleiotropy using either removal of outlier variants detected by MR-PRESSO outlier test or covariate adjustment in MultiPhenotype Mendelian Randomization^20^ (MPMR) in our simulations. After correction for either of these approaches, we no longer detected pleiotropy using the MR-PRESSO global test and minimization of inflation in the causal estimate due to pleiotropy was observed correspondingly (see **Supplementary Table 2**).

As a sensitivity check, we examined the effect of using overlapping samples in our two sample MR approaches, by simulating a one sample scenario where the effects of the variant on the exposure and the effects of the variant on the outcome are obtained from the same sample (see **Supplementary Table 3**). The effect of overlapping samples did not have any appreciable effect on the power for detecting pleiotropy or the accuracy of the causal estimates.

### Evaluation of pleiotropy for complex traits and diseases

Due to the observed inflated type I error rate of the Q test in our simulations, we focused specifically on MR-PRESSO and MR-Egger’s Intercept test in their application to summary-level GwA data. We applied MR-PRESSO and MR-Egger’s intercept test to all possible pairs of >80 complex traits and diseases retrieved from publicly available GWA datasets (see Table 2). In total, we conducted 4,250 tests for each of the four MR approaches. We accounted for multiple testing of the 4,250 tests using the Bonferroni correction. We note that this correction is overly stringent since many of the traits and diseases are correlated. Using a Bonferroni corrected threshold of *P*=1.17×10^-5^, the MR-PRESSO global test was statistically significant in 22 % (N = 922) of the 4,250 tests. Conversely, the MR-Egger intercept test was statistically significant in 0% (N=0) of the tests. When restricting to significant causal estimates in the IVW meta-analysis, we detected significance of the MR-PRESSO global test at a higher rate of 41% (N = 78) of the total number of tests (N = 191) whereas the MR-Egger test again was not significant in none of the tests. Very little overlap for significance was observed between the MR-PRESSO and MR-Egger intercept test, which can be explained by these tests being designed to detect different aspects of pleiotropy in MR (average pleiotropy for MR-Egger vs. outlier pleiotropy variants for MR-PRESSO).

**Table 2.** Comparison of two pleiotropy tests: MR-PRESSO global test and Egger Intercept.

**a) All pairs (5%)**
|  |  | Egger Intercept |  |
| --- | --- | --- | --- |
| | | P≥0.05 | P<0.05 |
| MR-PRESSO | P≥0.05 | 2280 (54%) | 151 (4%) |
|  | P<0.05 | 1663 (39%) | 156 (4%) |

**b) All pairs (BF)**
|  |  | Egger Intercept |  |
| --- | --- | --- | --- |
| | | P≥BF | P<BF |
| MR-PRESSO | P≥BF | 3328 (78%) | 0 (0%) |
|  | P<BF | 922 (22%) | 0 (0%) |

**c) Significant causal relationships (5%)**
|  |  | Egger Intercept |  |
| --- | --- | --- | --- |
| | | P≥0.05 | P<0.05 |
| MR-PRESSO | P≥0.05 | 73 (38%) | 25 (13%) |
|  | P<0.05 | 79 (41%) | 14 (7%) |

**d) Significant causal relationships (BF)**
|  |  | Egger Intercept |  |
| --- | --- | --- | --- |
| | | P≥BF | P<BF |
| MR-PRESSO | P≥BF | 113 (59%) | 0 (0%) |
|  | P<BF | 78 (41%) | 0 (0%) |

Comparison of the MR-PRESSO global test and the Egger test (student test of the intercept of the Egger regression) for a significance threshold of 5% (a, c) and for a Bonferroni threshold (BF) (b, d). Counts are either reported for all outcome/exposure pairs (a, b) or restricted to significant causal pairs (c, d) with a significant causal $P$ at the Bonferroni Threshold. BF: Bonferroni threshold.

### Control of pleiotropic effects through outlier removal

Next, we evaluated two strategies to correct for pleiotropic effects in MR tests (see Table 3), including: 1) removing outliers detected by the MR-PRESSO outlier test (at Bonferroni threshold); and 2) adjusting for significant covariates singly or all together from the main MR test (e.g. significant slope in IVW meta-analysis at Bonferroni threshold). We observed that the outlier removal approach using the MR-PRESSO outlier test was effective in eliminating statistical significance in the MR-PRESSO global test in 46% (N=422) of the 922 tests. Furthermore, the covariate adjustment approach-defined by accounting for traits that showed statistically significant results from the slope of the main MR test-eliminated significance in the mR-PRESSO global test in 22% (N = 20) of the 93 tests when adjusted singly. When adjusted for all significant covariates in the same model, the covariate adjustment approach eliminated significance in 34% (N = 22) of the 42 tests. Taken altogether, the correction strategies were successful in 47% (N=438) of the 922 tests total. We note that the covariate adjustment approach is limited in that it requires a priori knowledge on the trait responsible for the pleiotropic effect.

**Table 3.** Correction for pleiotropy in Mendelian randomization using two different approaches: MR-PRESSO outlier test and covariate adjustment.

|  | **MR-PRESSO**<br>**outlier removal** | **Single-**<br>**covariate**<br>**adjustment** | **All-**<br>**covariate**<br>**adjustment** |
| --- | --- | --- | --- |
| **Total corrected** | 422 | 20 | 22 |
| **Total remaining pleiotropy** | 500 | 73 | 42 |
| **Fraction corrected pleiotropy** | 0.46 | 0.22 | 0.34 |

We assessed 922 exposure-outcome pairs for which the pairwise MR-PRESSO test is statistically significant at the Bonferroni threshold. For each of these pairs, we applied two correction strategies to account for pleiotropy. The two correction strategies include a MPMR model where we adjusted for one covariate at a time or all together in the same model. To select the covariates, we considered all exposures with a significant causal effect for the outcome in question using the Bonferroni threshold. We included only covariates with a correlation coefficient $< 0.3$ with the tested exposure, the correlation coefficient being calculated using the effect sizes of the exposure and the covariate of the variants associated with the tested exposure. Pleiotropy was considered as corrected if at least one of the adjustments led to a *P-value* higher than the significance threshold. The table reports counts of pairs with corrected pleiotropy (corrected pleiotropy adjusted *P-value* from MR-PRESSO global test is not significant at Bonferroni threshold) and pairs with remaining pleiotropy (corrected pleiotropy adjusted *P-value* from MR-PRESSO global test is still significant at Bonferroni threshold).

### Pleiotropy confounds causal estimates in MR tests

We evaluated the extent to which outliers as detected by the MR-PRESSO outlier test can cause bias in the causal estimates from MR. Using the MR-PRESSO distortion test, we compared the causal estimate from the IVW meta-analysis before and after removal of outlier variants detected by the MR-PRESSO outlier test (see **Online Methods**). Using a Bonferroni threshold, we observed in 2.5% (N = 2) of significant causal estimates (N = 81 total)), a significant distortion of-93% and 35%. Since the Bonferroni correction is overly stringent, we considered the commonly-used nominal threshold of *P* < 0.05 that the majority of MR studies to date have used for statistical significance. A significant distortion was observed in almost 10% (N = 22) of the causal relationships (N = 229 total) with a distortion between-168% and 189% (see Figure 2).

**Figure 2.**
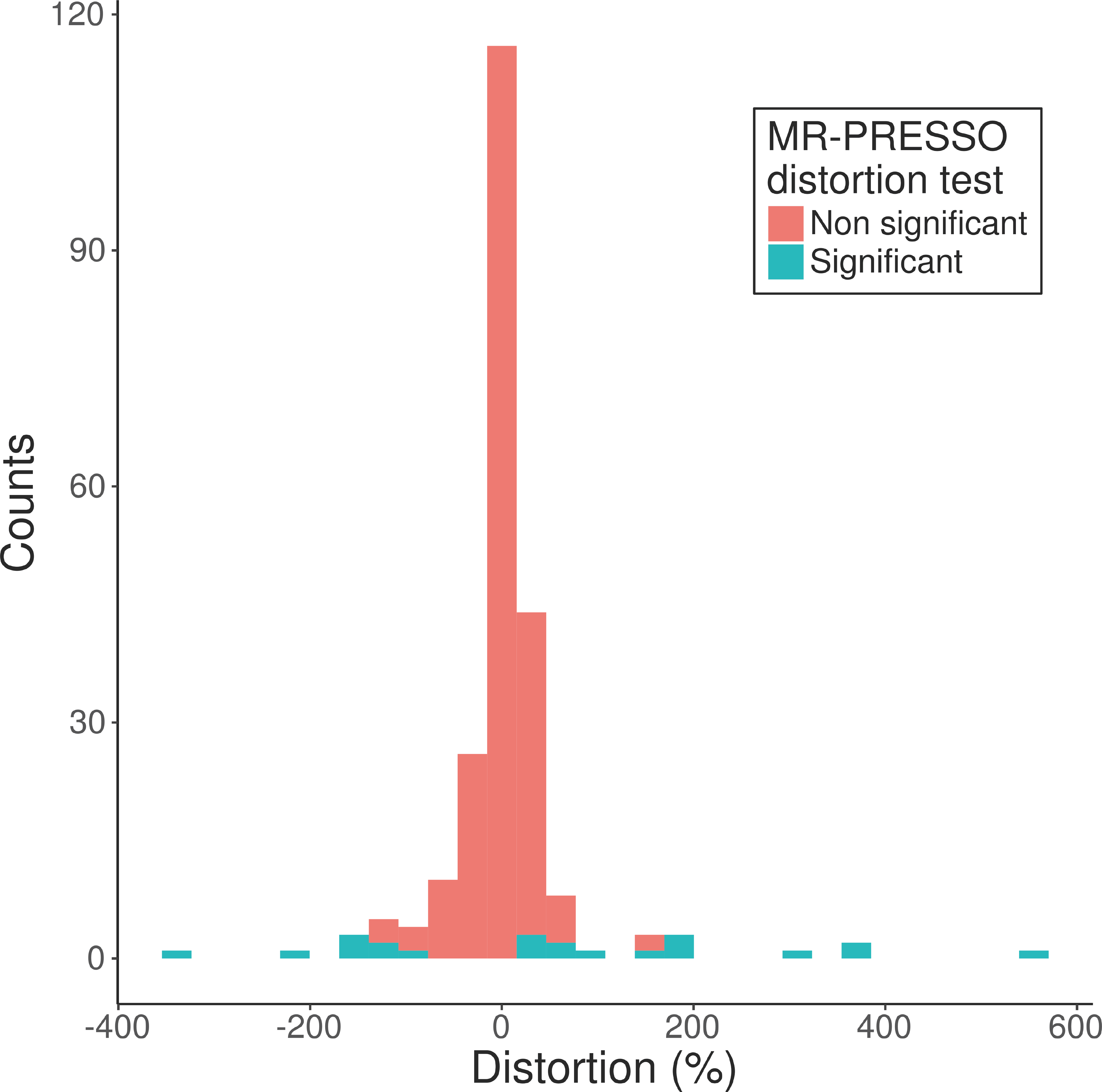
Distribution of the distortion of causal estimates before and after correction for pleiotropy using MR-PRESSO. The distortion coefficients are colored according to whether the distortion is statistically significant (blue) or not (red) at a threshold of *P* < 0.05 / 229 in the MR-PRESSO distortion test. The distortion estimate is defined as the percent change in the causal estimate as a result of outliers. It is defined as the (causal estimate from the IVW metaanalysis before removal of outlier variants minus causal estimate from the IVW metaanalysis after removal of outlier variants as detected by the MR-PRESSO outlier test) divided by the absolute value of the causal estimate from the IVW meta-analysis after removal of outlier variants as detected by the MR-PRESSO outlier test. A positive distortion represents a decrease in the outlier-corrected causal estimate whereas a negative distortion represents an increase in the outlier-corrected causal estimate.

As a representative example, we highlighted one relationship that exemplifies this type of confounding (see **Supplementary Figure 3**). We showed that the causal effect of body mass index (BMI) on C-reactive protein was estimated to be 0.39 (*P* = 7.02 × 10^-8^) by IVW. The MR-PreSsO global test showed statistical significance (*P* < 10^-6^) with it being driven by one outlier variant (rs2075650 in the *APOE* locus) whereas notably the MR-Egger intercept test was not significant (*P* = 0.92). Examining this further, we observed that this variant was highly pleiotropic with associations with several traits and diseases including Alzheimer’s Disease, body mass index, C-reactive protein, high-density lipoprotein cholesterol, low-density lipoprotein cholesterol, plasma triglycerides and waist circumference, hip circumference and waist-hip ratio at *P* = 6 x 10^-4^ (Bonferroni corrected *P* = 0.05 / 82; see **Supplementary Table 4**). Furthermore, this variant was associated with several other traits and diseases in the public NHGRI-EBI GWAS catalog^21^ (*P* < 5 × 10^-8^; see **Supplementary Table 5**). After removing this outlier variant, we observed a lower estimation of the causal estimate of BMI on C-reactive protein (β_causal_ = 0.35, *P* = 3.45 × 10^-16^) with this single variant alone causing a 12% distortion in the causal estimate.

**Figure 3.**
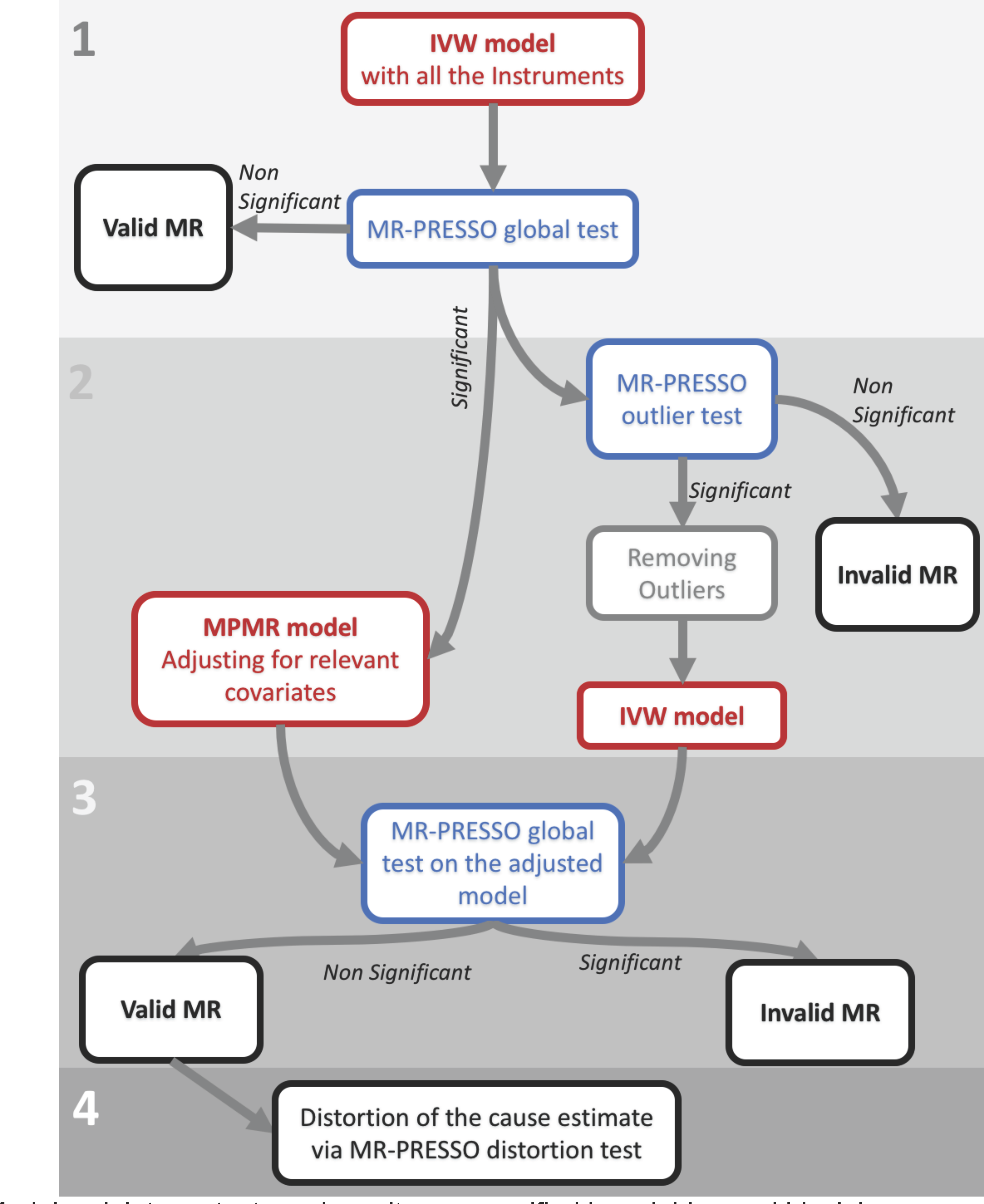
Flowchart of recommended steps for detection and correction of pleiotropy using the MR-PRESSO framework. Models, pleiotropy tests and results are specified in red, blue and black boxes respectively. The four distinct steps include: 1) initial MR analysis and test for pleiotropy using the MR-PRESSO global test; 2) adjustments for pleiotropy; 3) interpretation of the pleiotropy adjusted results; 4) assessment of significant differences in the causal estimate before and after removal of outliers using MR-PRESSO distortion test. Table 1: Evaluation of statistical power (%) at a significance threshold of 0.05 for different methods to detect pleiotropy in Mendelian randomization.

### Pleiotropy induces false positive causal relationships in a small proportion of MR tests

We evaluated the extent to which outliers as detected by the MR-PRESSO outlier test can confound causal estimates in MR tests in the most extreme form, by assessing how these outliers can induce “false positive causal relationships”, defined as those causal estimates that were no longer statistically significant in the outlier corrected IVW model but were previously significant in the naïve IVW model. Using this definition, we observed that 10% of false positive causal relationships (N = 24 out of 229 total tests) using the commonly used nominal P < 0.05 threshold and 1.2% (N = 1 out of 81 total tests) using the stringent Bonferroni corrected threshold (*P* < 1.17 × 10^-5^).

### Recommendation to correct for pleiotropy using MR-PRESSO

In Figure 3, we suggest the following flowchart to comprehensively detect and correct for pleiotropy using the MR-PRESSO framework. The steps include: 1) Perform the global MR-PRESSO test to assess whether pleiotropy is occurring between the biomarker and disease in the MR test; 2) If the global test is statistically significant, then adjust for known covariates in a MPMR model and re-test for pleiotropy with the global MR-PRESSO test; 3) If pleiotropy remains, perform the outlier test in MR-PRESSO to identify offending outlier pleiotropic variants; 3) Remove outlier pleiotropic variants from the MR test and re-test for pleiotropy with MR-PRESSO; 4) Perform test of significant differences in the causal estimate of the MR test, before and after pleiotropy correction; and 5) Use the pleiotropy corrected MR test as the causal estimate. We note that there are some scenarios where neither outlier removal or adjustment for relevant covariates will completely correct for pleiotropy in MR. In these instances, we believe that the causal estimates remain biased and should be interpreted with caution.

### Causal relationships inferred from MR testing

We identified 191 significant causal relationships out of a total number of 4,250 MR tests using the Bonferroni corrected *P* = 0.05 / 4,250 = 1 × 10^-5^. We note that many of the traits are closely related to each other (e.g. BMI/waist circumference/hip circumference, and low-density lipoprotein cholesterol (LDL-C)/total cholesterol) and hence the tests are not completely independent of each other. After correcting for pleiotropy via outlier removal using the MR-PRESSO outlier test, we validated known causal relationships including LDL-C on coronary artery disease (CAD) (β_causal_=0.52, *P*=5.15 ×10^-12^)^22^, systolic blood pressure on CAD (β_causal_=0.05, P=1.78 ×10^-6^)^23-25^, BMI on type 2 diabetes (β_causal_=0.76, *P*=2.19 ×10^-9^)^26^ and BMI on C-reactive protein (β_causal_=035, *P*=3.45x10^-16^), amongst the strongest findings. Furthermore, we observed an effect of BMI on uric acid (β_causal_=0.31, *P*=3.29×10^-15^)^27^ and plasma triglycerides (β_causal_=0.20, *P*=6.9 ×10^-15^)^12^, although these have significant pleiotropy even after correction via outlier removal using the MR-PRESSO outlier test.

## Discussion

In summary, we have evaluated pleiotropy in the context of MR testing across pairwise comparisons of a large number of complex traits and diseases. We have: i) developed the MR-PRESSO framework to detect and correct for pleiotropy in MR testing using GWA summary statistics; (ii) showed that pleiotropy occurs 41% amongst causal relationships inferred by MR between complex traits and diseases; (iii) observed distortion between-168% and 189% in the causal estimates of MR due to this pervasive pleiotropy; and (iv) showed that pleiotropy can be minimized and corrected in a large proportion of the cases using outlier detection (MR-PRESSO) and/or secondary phenotype adjustment (MPMR) of pleiotropic variants when the mediating pleiotropic trait is known.

In this study, we showed that the MR-PRESSO global test outperforms previous methods (MR-Egger and Q test) to detect pleiotropy in a MR framework. MR-Egger and Q test have inherent limitations. First, MR-Egger has been proposed as a method to assess pleiotropy on average across all instruments. However, it relies on the InSIDE (Instrument Strength Independent of Direct Effect) condition, which is untestable, and it has been noted that the adjustment for pleiotropy results in up to 30% reduction in statistical power in the causal estimate, which we confirmed by simulation^18^. Furthermore, the Q test can detect pleiotropy by testing whether a subset of variants have heterogeneous effects^28^. However, the Q test shows significant inflation in type I error in our simulations. The MR-PRESSO outlier test has the advantage in that it can identify those offending individual pleiotropic outlier variants for removal from any MR test. Importantly, MR-PRESSO can be used within the framework of the IVW metaanalysis and therefore retains statistical power of the approach, unlike MR-Egger regression.

By applying the MR-PRESSO global test to detect pleiotropy in a wide array of complex traits and diseases, we observed pleiotropy in approximately 41% of inferred casual relationships. This is consistent with emerging evidence that disease-associated variants identified from GWA studies have effects on other related traits^10^. Since these variants are used as instrumental variables in multiple instrument MR analysis, it’s likely that at least some number of these variants do not meet the ‘no pleiotropy assumption’ in MR. MR-PRESSO can detect those variants that do not meet this assumption and can be removed to maintain the validity of MR. Our results indicate pleiotropy is commonplace and highlight the need to make pleiotropic evaluation for variants acting as instrumental variables a necessary and standard test when performing MR analysis.

Pleiotropy in MR has direct implications for genetics-guided drug discovery and validation. Accurate estimates of causal estimates between a biomarker and disease can inform dose-response curves for drug efficacy and safety^29^ which are guided by the relationship of variants and their associations with the biomarker and disease. In the present study, we show that pleiotropy can induce bias by distorting the causal estimates in MR and that this bias is widespread. Secondly, there is increasing interest in using surrogate endpoints for drugs in clinical trials. Identifying true causal relationships using MR can guide those biomarkers that are causal and hence identify those surrogate endpoints that are most relevant to disease^30^.

The current study has several strengths. This approach is powerful because it works within the framework of IVW which has higher statistical power than other approaches such as MR-Egger. Furthermore, the MR-PRESSO global test is adequately powered to detect pleiotropy amongst even a small subset of loci; it has >75% to detect pleiotropy amongst 5% of variants with pleiotropic effect at the same level of causal effect. Finally, the MR-PRESSO framework is flexible and can be used in several different MR tests including IVW, MPMR and even within the framework of MR-Egger. The method also has limitations. There were a few instances where certain MR tests were shown to be statistically significant in the MR-PRESSO global test but the MR-PRESSO outlier test was not able to identify the outlier pleiotropic variants. This could be due to a subset of variants that have low-level pleiotropy where collectively the sum signifies a significant deviation from the null but each variants alone is not strong enough to be detectable by the MR-PRESSO outlier test. Furthermore, several GWA consortia utilize the same cohorts and study samples; therefore, some GWA summary statistics may have overlapping samples. In simulations, we evaluate how overlapping samples affect the power of our test across a range of tested scenarios. The power for detecting pleiotropy did not change substantially. Finally, because MR-PRESSO utilizes simulations; the processing time to apply the method can be slower compared to other methods.

In summary, we have developed and implemented a novel statistical framework called MR-PRESSO, to detect and correct for pleiotropy amongst variants used as instrumental variables in MR. By applying this framework to pairwise comparisons of a set of >80 complex traits and diseases, we show that pleiotropy is widespread, highlighting the need to correct for this pervasive bias in order to maintain the validity of causal inference testing in MR.

## Online Methods

### General assumptions of Mendelian randomization

MR utilizes genetic variants as the instrumental variable (IV) and estimates causal effects as ratio estimates between the genetic effect on the outcome and the genetic effect on the exposure. **Supplementary Figure 1** illustrates a standard MR framework (see **Supplementary Material** for further description of methods).

Our approach utilizes multiple IVs and extends on the framework of inverse variance-weighted, fixed effects meta-analysis (IVW meta-analysis). The IVW meta-analysis consists of fitting a weighted linear regression with fixed intercept of 0 between the set of effect sizes on the outcome and the effect sizes on the exposure, with the inverse of the variance of the effect sizes on the outcome as weights.

The validity of MR analysis relies on three assumptions (**Supplementary Figure 1**): 1) The variant (i.e. IV) is associated with the exposure; 2) The variant is independent of all confounders of the exposure-outcome relationship; 3) The variant is independent of the outcome conditioned on the exposure and all confounders of the exposure-outcome association (i.e. exclusion restriction). Violation of the third assumption, the exclusion restriction criterion, is a direct consequence of pleiotropy (**Supplementary Figure 2**).

### Limitations of existing methods to detect and correct for pleiotropy in MR

We evaluated pleiotropy in MR testing using three methods: MR-Egger regression, the Q test, and our newly developed method MR-PRESSO. The MR-Egger regression provides a test for average pleiotropy in multi-instrument MR (MR-Egger test)^18^. The intercept of the MR-Egger regression can be interpreted as the average pleiotropic effect across all IVs. The Q test is traditionally used to identify outliers and has been applied in the context of MR by Greco et al.^19^ to detect pleiotropy. Pleiotropy can induce heterogeneity of the individual ratio estimates. Therefore, the Q test can be used in the context of multi-instrument MR to test for over-dispersion.

Both of these methods have several limitations. MR-Egger assumes that a large proportion of the IVs is affected by pleiotropy for the intercept to capture the average pleiotropic effect. It also relies on the InSIDE (Instrument Strength Independent of Direct Effect) condition to hold, which is an untestable assumption. In addition, the estimation of an additional parameter, namely the intercept of the linear regression, substantially decreases the power of MR-Egger compared to IVW meta-analysis. Furthermore, the Q test only allows for detection of pleiotropy amongst the IVs; importantly, however, the Q test does not provide a way to correct for pleiotropy.

### MR-PRESSO: MR Pleiotropy RESidual Sum and Outlier

To overcome limitations of previous approaches, we proposed a novel framework called MR-PRESSO (MR Pleiotropy RESidual Sum and Outlier) for detecting and correcting pleiotropy in multiple IV MR using GWA summary statistic data.

The principle behind MR-PRESSO extends on the Q test (see Figure 1). IVW metaanalysis using multi-instrument MR can be performed using linear regression of the variant’s (IV) effect sizes of the exposure and outcome^17^. In the ideal scenario where pleiotropy does not exist (**Supplementary Figure 1**), all variants should reside on the regression line with the slope equal to the true causal effect of the exposure to the outcome. However, when a subset of variants are subject to pleiotropy (**Supplementary Figure 2**), these variants will deviate from the true slope of the regression line.

The MR Pleiotropy RESidual Sum and Outlier (MR-PRESSO) framework is comprised of three components: 1) detection of pleiotropy (and violation of the restriction criterion assumption, **Supplementary Figure 2**) in MR (global test); 2) correction by removal of offending IVs that exhibit pleiotropy (outlier test); and 3) testing of significant differences in the causal estimates before and after MR-PRESSO correction (distortion test).

For the MR-PRESSO global test, the approach is comprised of four steps: 1) for each variant *j*, we removed the specific variant in question and calculate the slope, denoted 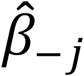, of the regression line on the remaining variants; 2) we calculated the observed residual sum of squares (RSS) as the difference between the observed effect size of the variant on the outcome (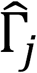) and the predicted effect size of the same variant on the outcome, calculated as the product of the slope (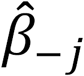; obtained in 1) and the effect size of the same variant on the exposure 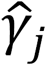, 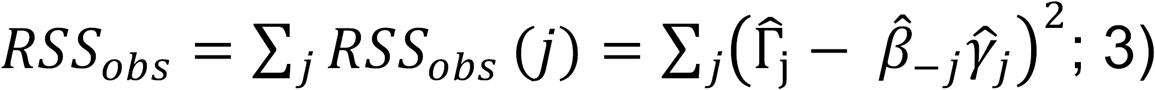 we calculated the expected RSS by drawing on the effect sizes on both the outcome 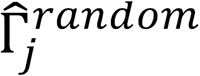 and exposure 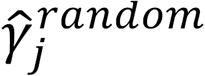 from Gaussian distributions using the standard errors of the variants on the exposure and outcome, respectively 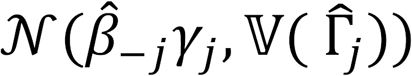 and 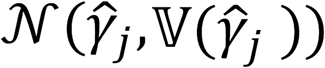. The procedure is repeated multiple times (K) to obtain a null distribution of the RSS, 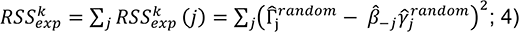 an empirical *P* can be computed as the number of expected RSSs greater than the observed RSS divided by the total number of times the procedure is repeated.

For the MR-PRESSO outlier test, we can identify outlier variants for removal of IVs with pleiotropy (offending variants) by utilizing the observed and expected distribution of the tested variant ftSS_obs_(j) and the K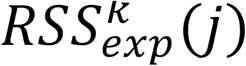 (Figure 1). We next computed a *P-value* per variant after a Bonferroni correction for the number of tested variants.

For the MR-PRESSO distortion test, we evaluated the statistical significance of the causal estimate before and after correction for pleiotropy via removal of outlier pleiotropic steps (from the outlier test of MR-PRESSO). To quantify this, we compared the original causal estimate 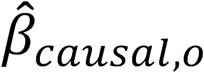,_0_ and the corrected causal estimate after removing outliers identified by MR-PRESSO 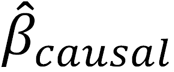. To compare the two estimates, we calculated the distortion as the percentage of the true causal estimate that is altered by pleiotropy. Mathematically, we define the distortion as

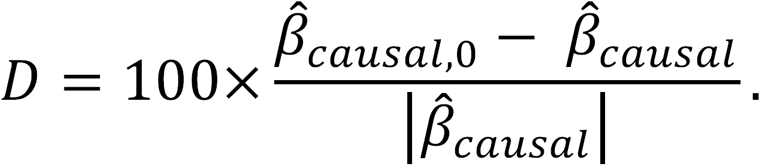

Normalizing by the absolute value of the corrected causal estimate allows to define a positive and negative distortion respectively as a decrease and increase in the corrected causal estimate regardless of the the signs of the causal estimates. To test for statistical significance of this value, we calculated an empirical *P-value* by generating a null distribution. The null distribution is generated by adding to the *n_o_* variants detected as outliers by MR-PRESSO, *n_E_*-2*n_o_* variants drawn with replacement from the set of non-outlier variants and maintained to *n_E_*-*n_o_*. We repeated this procedure 10,000 times to compute the null distribution of the distortion. An empirical *P-value* is then calculated as the number of times that NB is greater than the null estimates divided by 10,000. We have implemented the MR-PRESSO framework using the R Project for Statistical Computing (https://github.com/rondolab/MR-PRESSO).

### Multi-Phenotype Mendelian randomization: Correction for known pleiotropy

After evaluating the role that pleiotropy plays amongst a wide array of MR tests, we assessed various approaches to correct for this bias. Specifically, we examined the intercept in MR-Egger, MR-PRESSO outlier removal and MPMR. MR-Egger and MR-PREsSo were described above. We had previously developed a method called MPMR (Multi-Phenotype MR)^20^ which considers known confounders (E_2_ in **Supplementary Figure 2**) for the IV (Gj in **Supplementary Figure 1** and **2**) and outcome (Y in **Supplementary Figure 1** and **2**) and adjusts for these known confounders while performing MR analysis. In practice, MPMR is implemented by fitting a weighted linear regression by regressing 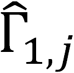 on 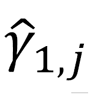 and 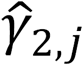. The 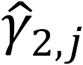 are the genetic effects of the variants on the known confounder E_2_. The model can be easily extended to include more confounders. In this study, a fixed intercept of 0 was used in all MPMR models. The advantage of MPMR is that it utilizes information on known confounders to control for bias whenever available. In the case where there are additional confounding effects between the pleiotropic effect mediator and the outcome of interest, residual mediated pleiotropic effect may occur even after adjusting for the mediator with MPMR.

### Simulation Framework

We performed simulations to evaluate the statistical properties (type I error and power) to detect pleiotropy, as well as the power of detecting causal effects under existing methods (Q test and MR-Egger) and MR-PRESSO. We simulated the standard MR framework shown in **Supplementary Figure 2b** with an outcome as well as two exposures Ei and E_2_. We induced pleiotropy in a subset of variants by associating with E_2_ either in a directional manner (effect sizes associated with E_2_ all positive) or in a balanced manner (effect sizes associated with E_2_ either positive or negative). We considered several scenarios using the following parameters of 1) causal effect E_1_ and E_2_ on the outcome; 2) type of pleiotropy: no pleiotropy, directional or balanced; 3) percentage of pleiotropic variants: 1, 5, or 10%; and 4) sample overlapping between the estimation of variant effects on the exposure and on the outcome.10,000 simulations were performed per scenario.

### Curation of genome-wide association (GWA) summary statistics

We retrieved publicly available genome-wide association (GWA) summary statistics data^31-66^ for 82 complex traits and diseases (see Supplementary Material). We performed the following steps to ensure that all datasets were uniform and standardized. For each, we retrieved the appropriate variant annotation (build, rsid, chromosome, position, reference and alternate alleles) and summary statistics (effect size, standard errors, *P-values* and sample size of the study). All variant coordinates (chr, pos) were lifted over to hg19 using the UCSC Genome Browser LiftOver Tool. We imputed Z-scores of variants using ImpG^67^ using 1000 Genomes Phase 3 European panel^68^ (N=503) as a reference panel. Effect sizes, standard errors and *P-values* were then calculated using the variance of the trait estimated from genotyped variants and allele frequencies calculated on the same subset of individuals from the 1000 Genomes reference panel. Sets of GWA-significant variants were manually retrieved from the corresponding GWA manuscripts. In total, we retrieved GWA summary statistics for 82 traits and diseases (see **Supplementary Table 6**).

### Detection and correction of pleiotropy using MR-PRESSO or covariate adjustment in MPMR

We applied MR-PRESSO to all possible exposure-outcome pairs of 82 traits and diseases, and then compared the results of this test to those obtained from MR-Egger. In total, we performed 4,250 MR tests. We compared these results to other MR pleiotropy approaches including MR-Egger and the Q test. Next, we evaluated two strategies to correct for significant pleiotropy detected from our MR-PRESSO test. First, we included covariates in our Multi-Phenotype Mendelian randomization (MPMR) model, either one by one. We considered only covariates with a statistically significant causal effect (slope of the IVW meta-analysis using a Bonferroni cut-off. Furthermore, to account for co-linearity, we included only covariates with a correlation coefficient < 0.3. Second, we corrected for pleiotropy using MR-PRESSO by removing offending variants that are statistically significant from the slope. 1,000,000 simulations to calculate the empirical *P-values* were performed for the MR-PRESSO global and outlier tests. 10,000 simulations were computed to calculate the empirical *P-values* for the MR-PRESSO distortion test.

## Acknowledgements

We thank the various genome-wide association consortia for generously sharing the genome-wide association summary statistics. R.D. was supported by an American Heart Association Cardiovascular Genome-Phenome Discovery grant (15CVGPSD27130014). B.N. is supported by 1R01MH094469 and 1R01MH107649-01 from the National Institutes of Health.

## Author contributions

M.V. and C-Y.C. contributed to study conception, data analysis, interpretation of the results and drafting of the manuscript. R.D. and B.N. contributed to study conception, interpretation of the results and critical revision of the manuscript.

